# Pannexin 2 restrains ER stress-induced Ca^2+^ dysregulation and inflammatory cardiomyocyte death

**DOI:** 10.64898/2026.01.27.702168

**Authors:** Anusua Sarkar, Li Zhu, Xiaoguang Margaret Liu, Lufang Zhou

**Affiliations:** Departments of Surgery, The Ohio State University; Biomedical Engineering, The Ohio State University; Chemical and Biomolecular Engineering, The Ohio State University; Davis Heart & Lung Research Institute, College of Medicine, The Ohio State University

**Keywords:** pannexin 2, ER stress, calcium homeostasis, pyroptosis

## Abstract

**Background:** Endoplasmic reticulum (ER) stress and ER-mitochondria Ca^2+^ dysregulation contribute to cardiomyocyte injury, yet endogenous regulators at ER-mitochondria interfaces that restrain this cascade remain poorly defined. Pannexin 2 (Panx2), the most structurally divergent pannexin isoform, has been implicated in stress response, but its cardiac localization and function are unclear.

**Methods:** Panx2 localization and function were assessed in human AC16 cardiomyocytes using high-resolution confocal imaging and complementary loss- and gain-of-function approaches during thapsigargin-induced ER stress, with validation in adult mouse ventricular cardiomyocytes.

**Results:** Panx2 localizes predominantly to the ER and mitochondria-associated membranes, rather than the plasma membrane. Panx2 knockdown reduced ER Ca^2+^ stores and increased basal cytosolic and mitochondrial Ca^2+^. During ER stress, Panx2 deficiency markedly amplified Ca^2+^ dysregulation, mitochondrial dysfunction, unfolded protein response activation, and cytotoxicity, with PERK-dominant signaling and increased IRE1a activation. Notably, PERK inhibition preferentially rescued the Panx2-deficient phenotype, providing the greatest improvement in viability and reduction in cytotoxicity. Panx2 deficiency also enhanced inflammasome/ pyroptotic signaling via the NLRP3-caspase-1-gasdermin D axis. Conversely, Panx2 overexpression suppressed PERK activation and attenuated ER stress-induced injury. Panx2 ablation similarly sensitizes adult ventricular cardiomyocytes to ER stress.

**Conclusions:** Panx2 functions as an organelle-associated checkpoint at ER-mitochondria interfaces that stabilizes Ca^2+^ homeostasis and limits PERK-dominant ER stress signaling and inflammatory cell death programs in cardiomyocytes, providing a mechanistic framework for cardiomyocyte loss in cardiac disease.

**Research Perspective:** *1. What New Question Does This Study Raise?:* Does Panx2 serve as an endogenous “stress threshold” determinant in cardiomyocytes in vivo, governing when ER stress transitions from adaptive signaling to PERK-driven mitochondrial failure and inflammasome-associated inflammatory cell death during cardiac injury (e.g., ischemia-reperfusion, pressure overload, or cardiometabolic stress)?

*2. What Question Should Be Addressed Next?:* In clinically relevant models of heart disease (ischemia-reperfusion), test whether cardiomyocyte-specific Panx2 loss or augmentation alters infarct size, arrhythmia burden, ventricular remodeling, and functional recovery, and determine whether targeting the Panx2-PERK axis (e.g., selective PERK modulation in the acute reperfusion window or Panx2-directed strategies) reduces cardiomyocyte loss without impairing adaptive stress signaling needed for repair.

## Introduction

Cardiomyocyte loss is a central determinant of adverse remodeling and heart failure after myocardial infarction and ischemia-reperfusion injury^1, 2^. Endoplasmic reticulum (ER) stress and maladaptive unfolded protein response (UPR) signaling, coupled to ER-mitochondria Ca²⁺ transfer, can trigger mitochondrial dysfunction and regulated inflammatory cell death programs in the stressed myocardium^3–6^. However, the endogenous regulators at ER-mitochondria interfaces that stabilize Ca²⁺ homeostasis and prevent escalation from adaptive stress signaling to cardiomyocyte death remain incompletely defined.

Pannexins are large pore, single-membrane hemi-channels that can mediate fluxes of ions and small signaling molecules (e.g., ATP, amino acids, and other metabolites), influencing paracrine signaling and cellular stress responses^7–9^. Among the three isoforms (Panx1-3), Panx1 is the most extensively studied and has been implicated in inflammation, metabolic regulation, and ionic homeostasis^10–14^. In contrast, the biological functions of pannexin 2 (Panx2), the largest, most structurally divergent pannexin, remain much less defined, especially in adult peripheral tissues. Although initially considered central nervous system-enriched and functionally redundant with Panx1^8, 15–17^, recent transcriptomic and proteomic datasets support Panx2 expression in a broad range of non-neuronal tissues, including the heart^18–20^.

Recent structural studies indicate that Panx2 assembles as a heptameric channel, distinct from the hexameric architecture of Panx1 and connexins^21, 22^. Notably, Panx1 and Panx2 share low sequence identity (∼16%), highlighting their distinct evolutionary trajectories and suggesting isoform-specific regulation and function. Emerging evidence further links Panx2 to cellular stress outcomes context-dependent manner^19, 23, 24^. Importantly, recent endogenous and high-resolution imaging studies in diverse cell types localize Panx2 primarily to intracellular compartments rather than the plasma membrane, with enrichment at the ER, trans-Golgi network, and/or mitochondria-associated membranes (MAMs)^25–27^. These specialized microdomains coordinate Ca²⁺ handling, bioenergetics, and stress signaling, processes central to cardiomyocyte survival under pathological stress^28^. Despite these compelling biochemical, functional, and structural insights, Panx2 expression, subcellular distribution, and functional contribution to ER stress responses in cardiomyocytes have not been systematically defined.

Here, we address this critical gap by defining the expression and localization of Panx2 in human AC16 cardiomyocytes and by determining its role in ER stress responses. Using complementary gain- and loss-of-function approaches, ER stress paradigms, and primary adult mouse cardiomyocytes, we tested the hypothesis that Panx2 stabilizes ER Ca^2+^ homeostasis and limits maladaptive ER stress signaling. Our results demonstrate predominant intracellular/MAM-associated Panx2 localization in cardiomyocytes and show that Panx2 deficiency amplifies Ca^2+^ dysregulation, PERK-dominant UPR activation, mitochondrial dysfunction, and inflammasome-associated pyroptotic cell death. These results establish Panx2 as a novel regulator of cardiomyocyte stress adaptation and provide a conceptual framework for further defining its roles in cardiac physiology and disease.

## Methods and Materials

### Animals

All animal procedures adhered to the *Guide for the Care and Use of Laboratory Animals* (NIH Publication No. 85-23, revised 2011) and were approved by the Institutional Animal Care and Use Committee (IACUC) at Ohio State University (OSU). Adult C57BL/6J mice were purchased from Jackson Laboratories and maintained in the OSU animal facility under standard housing conditions. Panx2 knockout (Panx2^−/−^) mice were generated from a targeted tm1a knockout-first allele obtained through the UC Davis Knockout Mouse Project (KOMP) Repository (Panx2tm1a(KOMP)Wtsi) and subsequently bred with mice expressing a ubiquitous CMV-driven Cre recombinase to produce constitutive Panx2-null animals.

### Cell culture

AC16 cells were obtained from ATCC and maintained in DMEM/F12 (Gibco) supplemented with 2 mM L-Glutamine, 12.5% fetal bovine serum (FBS), and 1x penicillin-streptomycin (P/S). HEK 293A cells were cultured in DMEM (Gibco) containing 2 mM L-Glutamine, 10% FBS, 1x P/S, and 1 x MEM non-essential amino acids. Cells were incubated at 37 °C in a humid atmosphere with 5% CO_2_ and maintained between 10%-80% confluence in T25 or T75 flasks. Cell density and viability were measured using either a TC20 Automated Cell Counter (Bio-Rad, Cambridge, MA, USA) or a hemocytometer with trypan blue exclusion (Fischer Scientific, Waltham, MA, USA).

### siRNA transfection

AC16 cells were seeded in 6-well plates and grown to ∼70% confluency before transfection. To knockdown Panx2 gene, cells were transfected with human *Panx2* siRNA vector (Santa Cruz Biotechnology, sc-106351) using Lipofectamine™ RNAiMAX (Life Technologies, Carlsbad, CA, USA) according to the manufacturer’s instructions. A mock-transfected group, treated with transfection reagent alone (i.e., without siRNA), served as the control. Cells were used for downstream experiments 24 h after transfection. Knockdown efficiency was confirmed by RT-qPCR and Western blotting.

### Adenovirus production and transduction

The human Panx2 gene was synthesized (Addgene, Watertown, MA, USA), PCR-amplified, and cloned into pENTR-D-TOPO vector (Life Technologies, K2400) to generate the entry vector pEntr-CMV-Panx2^29, 30^. The construction was sequence-verified using M13 forward and reverse primers. The adenoviral expression vector pAd-CMV-Panx2 was produced by LR recombination with the ViraPower™ Adenoviral Gateway™ Expression Kit (Life Technologies, K4930). For viral production, pAd-CMV-Panx2 was linearized with PacI, purified, and transfected into HEK293A cells using Lipofectamine 3000 (Invitrogen). Cells were harvested when >80% cytopathic effect was observed, followed by three freeze-thaw cycles to release crude viruses. High-tier stocks were amplified by infecting HEK293A cells at an MOI of ∼5 and collected 2-3 days post-infection when cells detached. Titers were determined using the QuickTiter™ Adenovirus Titer Immunoassay Kit (Cell Biolabs, San Diego, CA, USA). For transduction, AC16 cells at 60-70% confluence were infected with Panx2 adenovirus, and efficiency was confirmed by RT-qPCR and western blotting 48 h later.

### Western blotting

Cells were lysed in RIPA buffer containing protease and phosphatase inhibitor cocktails, followed by brief sonication and centrifugation at 21,000 x g for 20 min at 4 °C. Protein concentration in the clarified lysates was determined using a BCA assay. Equal amounts of protein (20-70 mg per lane) were separated by SDS-PAGE on NUPAGE 4-12% Bis-Tris gels (Life Technologies), transferred to PVDF membranes, and blocked for 1 h at room temperature with 5% non-fat dry milk in TBST (TBS+0.1%Tween 20). Membranes were incubated with primary antibodies overnight at 4 °C, washed in TBST, and then incubated with HRP-conjugated secondary antibodies (anti-rabbit or anti-mouse, as appropriate) diluted 1:2000 in 3% non-fat dry milk for 1 h at room temperature. Immunoreactive bands were visualized using Immobilon ECL Ultra (Millipore, Burlington, MA, USA) or SuperSignal West Atto (Thermo Fisher) and imaged on an Azure Biosystems C300 system (Azure Biosystems, Dublin, CA, USA).

### RNA isolation and transcriptional analysis

Total RNA was extracted from AC16 cells using TRIzol reagent, and cDNA was synthesized from 1 mg RNA using the High-Capacity RNA-to-cDNA™ Kit (Applied Biosystems). RT-qPCR was performed using SYBR Green Supermix (Bio-Rad) on a Cielo 3 (AIQ030) real-time PCR system (Azure Biosystems)^31^. The expression of genes was normalized to *GAPDH,* and relative expression was calculated using the 2^−ΔΔCt^ method. PCR primers are listed in supplemental Tables 1 and 2, and their specificity was verified by melt-curve analysis and 1% agarose gel electrophoresis.

### Cell viability and cytotoxicity assays

Cell viability was assessed using the MTT assay. AC16 cells (1×10^4^ cells/well) were seeded in 96-well plates and incubated overnight at 37 °C. Following treatment, 10 mL of MTT reagent (5 mg/mL stock) was added to each well along with 100 mL growth medium, and plates were incubated for 4 h at 37 °C. The medium was then replaced with 50 mL DMSO, and plates were incubated for 10 min to solubilize formazan crystals. Absorbance was read at 570 nm on a SpectraMax iD3 plate reader (Molecular Devices, San Jose, CA, USA).

Cytotoxicity was assessed using the CyQUANT™ LDH Cytotoxicity Assay Kit (Life Technologies) as previously described^32^. At the end of treatment, 200 mL culture supernatant was collected and transferred to a 96-well plate. Next, 100 < l of the LDH reaction mixture was added, followed by 30 min incubation at room temperature in the dark. Absorbance was measured at 490 nm with background correction at 680 nm using a SpectraMax iD3 plate reader. LDH release and percent cytotoxicity were calculated per the manufacturer’s instructions using background-corrected values.

### Adult mouse cardiomyocyte isolation and in-vitro experiments

Pannexins are large pore, single-membrane hemi-channels that facilitate the passage of ions and small signaling molecules (e.g., ATP, amino acids, metabolites), thereby regulating paracrine communication and stress responses^7–9^. Of the three isoforms (Panx1-3), Panx1 is best characterized and linked to purinergic signaling, inflammation, and ionic homeostasis in diverse tissues^10–14^. In contrast, Panx2, the largest and most structurally divergent pannexin, remains much less studied, particularly in adult peripheral organs. Although initially thought to be restricted to the central nervous system and functionally redundant with Panx1^8, 15–17^, transcriptomic and proteomic datasets now support robust Panx2 expression in multiple non-neuronal tissues, including the heart^18–20^. These observations suggest broader physiological roles for Panx2 than previously appreciated.

### Confocal imaging

#### Colocalization analysis

For ER localization, AC16 cells were seeded on 15-mm glass-bottom dishes and co-transduced with Panx2-dTomato adenovirus and an ER-targeted eYFP adenovirus. For mitochondrial localization, Panx2-expressing cells were stained with MitoTracker Green (100 nM). Plasma membrane localization was assessed by wheat germ agglutinin (WGA) staining, performed on Panx2-transduced AC16 cells as previously described^30^. Cells were imaged on a Leica Stellaris 5 confocal microscope (Teaneck, NJ, USA) using 561 nm laser excitation for dTomato and 488 nm excitation for ER-eYFP, MitoTracker Green, or WGA. Images were processed offline in ImageJ (NIH) for colocalization analysis.

#### Mitochondrial membrane potential and ROS

AC16 cells were loaded with MitoSox Red (500 nM, Invitrogen) and MitoView 633 (100 nM, Biotium)^33^ for 15 min at 37 °C in phenol-red-free medium. Fluorescence was captured using 405 nm (MitoSOX) and 638 nm (MitoView 633) laser lines.

#### Calcium dynamics

ER luminal Ca^2+^ was measured with Mag-Fluo-4 AM as described^34^. Briefly, following dye loading and a 30 min de-esterification, cells were transferred to Mag^2+^-free medium with 220 nM free Ca^2+^ after 1.5 mM MgATP addition. Cytosolic Ca^2+^ was assessed with Fluo-4 AM^35^. Both Mag-Fluo-4 and Fluo-4 fluorescence were acquired using the 488 nm laser line. Mitochondrial Ca^2+^ was measured using a mitochondrial-targeted RCamp genetic probe^36^. For RCamp imaging, cells were transduced with RCamp adenovirus and imaged 48 h post-transduction using the 561 nm laser line.

#### Calcein ethidium homodimer viability assay

Isolated adult cardiomyocytes were incubated with Calcein AM (2 mM) and Ethidium Homodimer-1 (EthD-1, 4 mM) for 30 minutes at 37 °C in phenol-red-free medium. After incubation, cells were gently washed and imaged immediately. Fluorescence was acquired using the 488 nm laser line for Calcein (live cells) and the 561 nm laser line for EthD-1 (dead cells). Cell viability was quantified as the percentage of Calcein-positive cells relative to the total number of Calcein-positive plus EthD-1-positive cells within each imaging field.

### Antibodies and chemicals

Primary antibodies for Western blots were purchased from Cell Signaling Technology (PERK, C33E10; p-PERK, 3179; IRE1a, 3294; GSMD, 39754; cleaved GSMD, 36425; NLRP3, 15101; Caspase-1, 3866; cleaved Caspase-1, 4199; GAPDH, 2118), Novus biologicals (p-IRE1a, NB100-2323), and Invitrogen (PANX2, 42-2900), respectively. Secondary antibody mouse Anti-Rabbit IgG was obtained from Cell Signaling Technology (93702). PVDF membrane was obtained from Thermo Scientific. Chemical inhibitors for ATF6a (Ceapin-A7), IRE1a (MKC-3946), and PERK (GSK2656157) were purchased from Cayman Chemicals. All other reagents were acquired from Sigma-Aldrich.

### Statistics

Data were presented as box and whisker plots unless otherwise specified. Statistical comparisons were performed using t-test, one-way ANOVA with Holm-Šidák post-hoc testing or two-way ANOVA with Tukey’s multiple comparisons test (GraphPad Prism). Normality was assessed using the Shapiro-Wilk test. P < 0.05 was considered statistically significant.

## Results

### Panx2 is predominantly intracellular and supports ER stress adaptation in AC16 cardiomyocytes

To define the subcellular localization of Panx2 in cardiomyocytes, AC16 cells were transduced with Panx2-dTomato adenovirus and stained with WGA to label the plasma membrane. Confocal imaging revealed that Panx2 displayed a predominantly intracellular distribution with no evident colocalization with WGA (Figure 1A). Instead, co-expression of Panx2-dTomato with an ER-targeted YFP marker showed extensive overlap between Panx2 and the ER network (Manders coefficient ∼0.97; Figure 1B). In addition, co-staining with MitoTracker demonstrated close spatial proximity between Panx2 and mitochondria (Figure 1C). These data demonstrate that Panx2 is predominantly intracellular and enriched at ER-mitochondria interfaces in AC16 cardiomyocytes.

**Figure 1.**
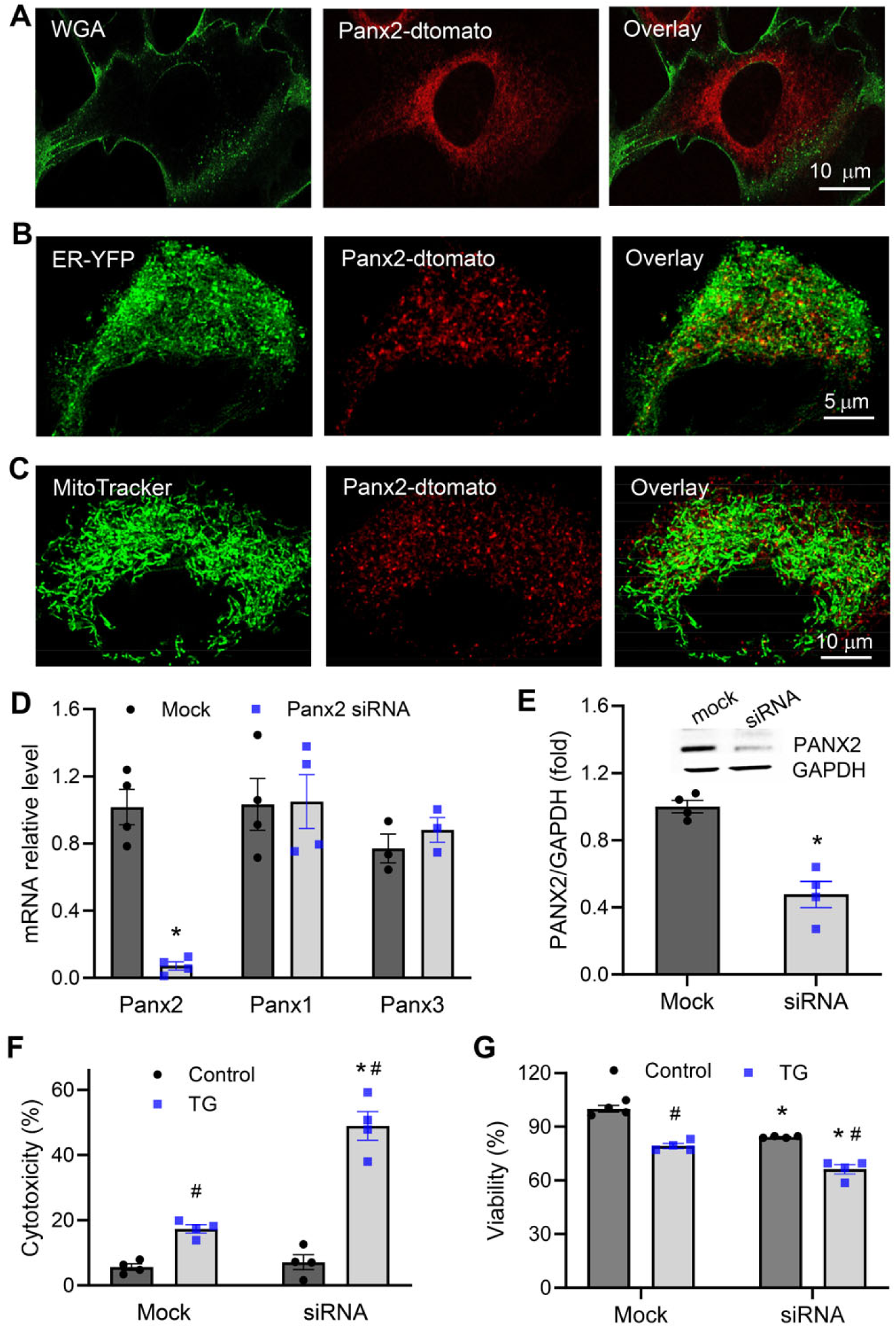
Panx2 is enriched at ER-mitochondria interfaces and regulates ER stress response in AC16 cardiomyocytes. (A) Representative confocal images of AC16 cells expressing Panx2-dTomato and stained with wheat germ agglutinin (WGA) to label the plasma membrane. Panx2 shows a predominantly intracellular distribution with minimal overlap with WGA. (B) Confocal images of AC16 cells co-expressing Panx2-dTomato and an ER-targeted YFP marker, demonstrating extensive overlap between Panx2 and the ER network. (C) Confocal imaging of Panx2-dTomato-expressing AC16 cells with MitoTracker staining, showing close spatial proximity between Panx2 and mitochondria. (D) RT-qPCR analysis of Panx2, Panx1, and Panx3 mRNA expression in AC16 cells following siRNA-mediated Panx2 knockdown. (E) Representative immunoblots and quantification confirming efficient depletion of Panx2 protein following siRNA transfection. (F) LDH release from mock- and Panx2 siRNA-transfected AC16 cells following TG (250 nM, 24 h) exposure. (G) MTT assay of cell viability in mock- and Panx2-deficient cells under basal conditions and post TG exposure. Data are presented as mean ± SEM. Statistical significance was determined by t-test (D-E) or two-way ANOVA (F-G). *: P < 0.05 vs. mock. #: P < 0.05 vs. control. n=4/group.

To assess the role of Panx2 in regulating cardiomyocyte stress response, AC16 cells were transfected with Panx2 siRNA. RT-qPCR detected a ∼90% reduction in Panx2 mRNA, while Panx1 and Panx3 transcript levels were unchanged (Figure 1D), demonstrating effective and specific *Panx2* knockdown. Efficient depletion at the protein level was confirmed by immunoblotting (Figure 1E). Following knockdown, cells were exposed to TG (250 nM) for 24 h to induce ER stress. LDH release assays showed that TG significantly increased cytotoxicity in mock-transfected cells and that Panx2 knockdown further increased TG-induced LDH release (Figure 1F). Consistently, MTT assays revealed reduced cell viability following TG treatment, with a significantly greater decline observed in Panx2-deficient cells (Figure 1G). It is worth noting that under basal conditions, Panx2 knockdown resulted in a modest but significant reduction in MTT signal without a corresponding increase in LDH release (Figure 1F-G), suggesting sublethal impairment in the absence of overt membrane damage.

### Panx2 loss disrupts ER, cytosolic and mitochondrial Ca^2+^ homeostasis

Because TG induces ER stress by inhibiting the SERCA and depleting ER Ca^2+^ store, we examined the impacts of Panx2 loss on Ca^2+^ homeostasis in TG-treated cardiomyocytes. ER luminal Ca^2+^ levels were measured using the low-affinity Ca^2+^ indicator Mag-Fluo-4 under basal conditions (control) and following TG exposure (6 h). TG markedly reduced ER Ca^2+^ levels in mock-transfected cells, with a significantly greater reduction observed in Panx2-deficient cells (Figure 2A). Under basal conditions, ER Ca^2+^ content was also significantly lower in Panx2 knockdown cells compared with mock-transfected cells (Figure 2A).

**Figure 2.**
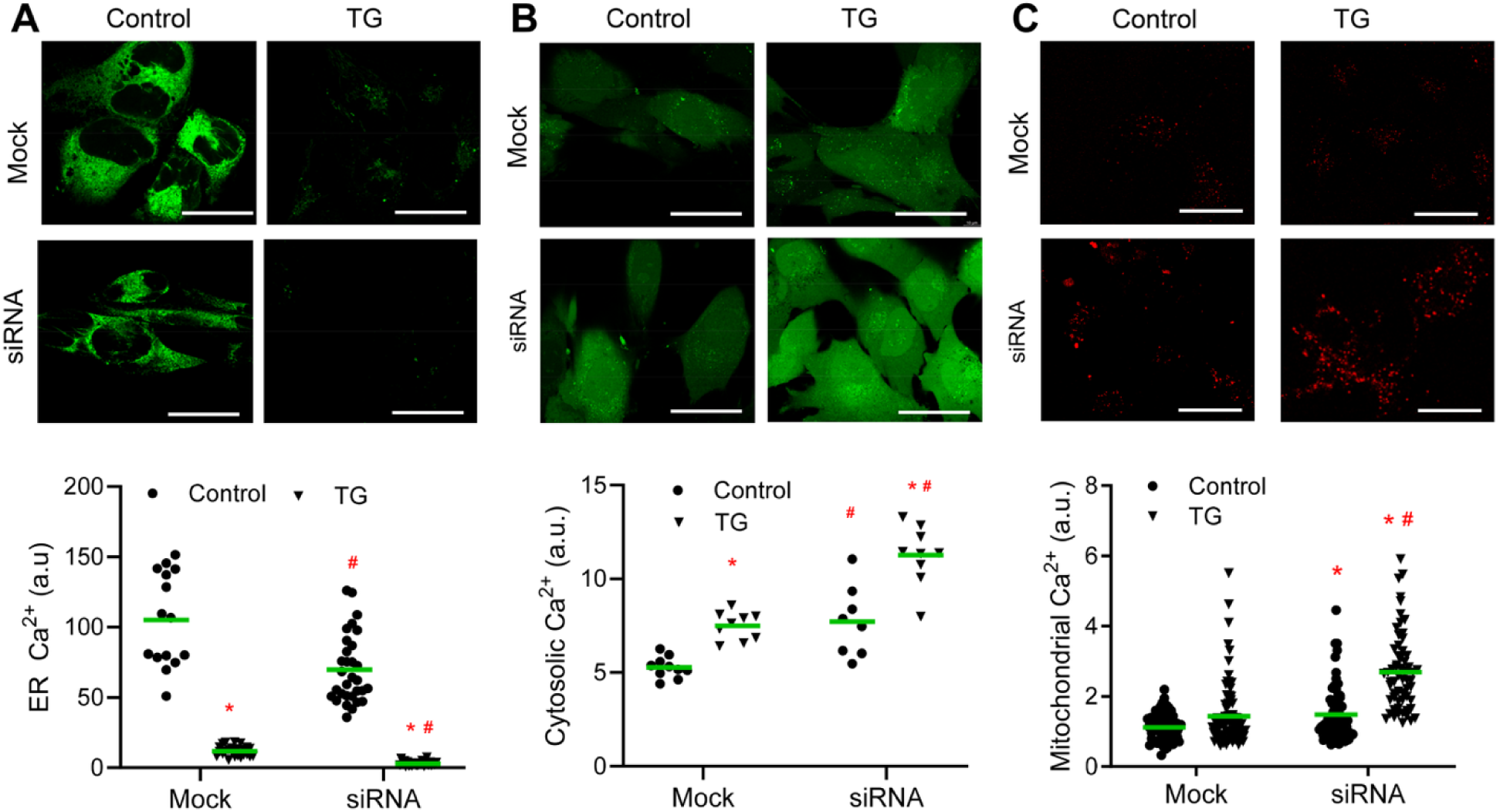
Panx2 knockdown alters intracellular Ca²⁺ homeostasis in AC16 cardiomyocytes. (A) Representative confocal images and quantification of ER luminal Ca^2+^ levels measured using Mag-Fluo-4 AM fluorescent dye. (B) Representative confocal images and quantification of cytosolic Ca^2+^ levels measured using Fluo-4 AM fluorescent dye. (C) Representative confocal images and quantification of mitochondrial Ca^2+^ levels measured using a mitochondrial-targeted RCamp genetic probe. Data are presented as mean ± SEM. Statistical significance was determined by two-way ANOVA. *: P < 0.05 vs. control. #: P < 0.05 vs. mock. Up to 40 cells from 4 different cultures per group were examined. Scale bars: 30 mm.

To assess downstream consequences of impaired ER Ca^2+^ handling, cytosolic and mitochondrial Ca^2+^ levels were also assessed. At baseline, both cytosolic (Figure 2B) and mitochondrial (Figure 2C) Ca^2+^ levels were significantly elevated in Panx2-deficient cells compared with mock-transfected controls. Following TG treatment, cytosolic Ca^2+^ levels increased significantly in mock cells, consistent with ER Ca^2+^ depletion following SERCA inhibition, whereas mitochondrial Ca^2+^ elevation did not reach significance (Figure 2B-C). In contrast, TG exposure resulted in remarkable increases in both cytosolic and mitochondrial Ca^2+^ in Panx2-deficient cells (Figure 2B-C). The magnitude of TG-induced Ca^2+^ elevation in both compartments was significantly greater in Panx2 knockdown cells than in mock-transfected cells.

### Panx2 deficiency amplifies UPR activation with PERK-dominant engagement

Given the marked disruption of Ca^2+^ homeostasis observed in Panx2-deficient cells, we next examined the impacts of Panx2 loss on ER stress signaling and the unfolded protein response (UPR) activation during a TG time course (baseline, 15, 30, and 60 minutes). In mock-transfected cells, TG increased phosphorylation of PERK (p-PERK/total PERK) and IRE1a (p-IRE1a/total IRE1α) and induced ATF6 cleavage by 60 minutes relative to baseline (Figure 3A-C). Panx2 knockdown cells exhibited significantly elevated p-PERK levels at baseline compared to mock cells and a further time-dependent increase following TG exposure, with p-PERK/PERK ratios reaching ∼5.8-fold above baseline at 60 minutes (Figure 4A). Similarly, TG induced robust IRE1a phosphorylation in both groups, with p-IRE1a levels consistently higher in Panx2-deficient cells at all time points examined (Figure 3B). Additionally, cleaved ATF6 levels were increased in Panx2 knockdown cells under basal conditions and further increased following TG exposure, whereas the total ATF6a remained unchanged (Figure 3C).

**Figure 3.**
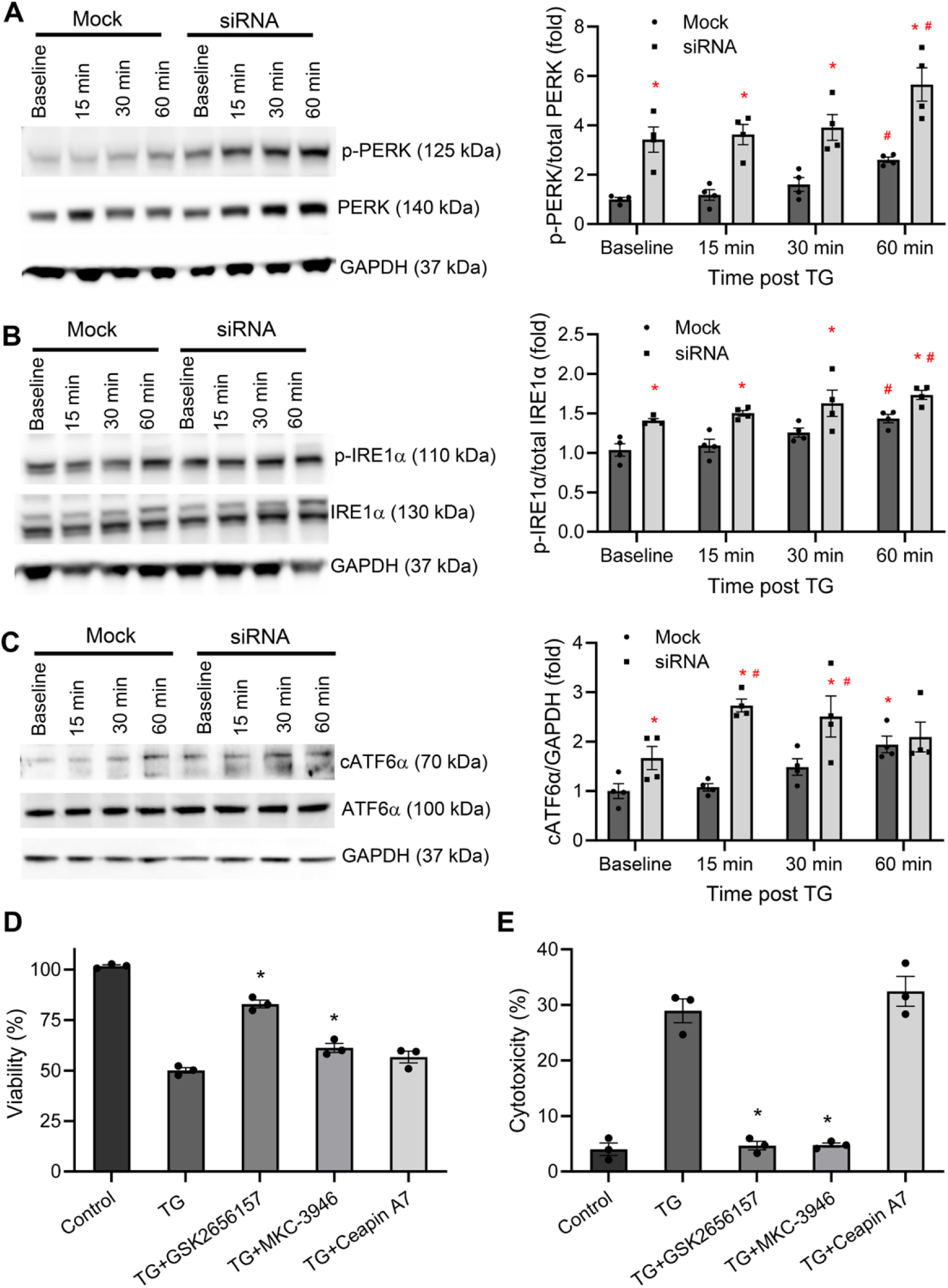
Panx2 deficiency amplifies TG-induced ER stress signaling. (A-C) Representative immunoblots and quantification of unfolded protein response (UPR) activation in mock- and Panx2 siRNA-transfected AC16 cells exposed to TG (250 nM) for 0, 15, 30, or 60 minutes. (A) Phosphorylated PERK (p-PERK) normalized to total PERK. (B) Phosphorylated IRE1a (p-IRE1a) normalized to total IRE1a. (C) Cleaved ATF6a. (D) MTT assay assessing viability of Panx2 siRNA-transfected AC16 cells treated with TG (250 nM, 24 h) in the presence or absence of PERK (GSK2656157, 0.3 mM), IRE1a (MKC-3946, 10 mM), or ATF6a (Ceapin-A7, 1 mM) inhibitors. (E) LDH release assay under the same conditions as in (D). Data are presented as mean ± SEM. Statistical significance was determined by two-way ANOVA. *: P < 0.05 vs. mock in A-C or TG in D-E. #: P < 0.05 vs. baseline. n=4/group/time point.

**Figure 4.**
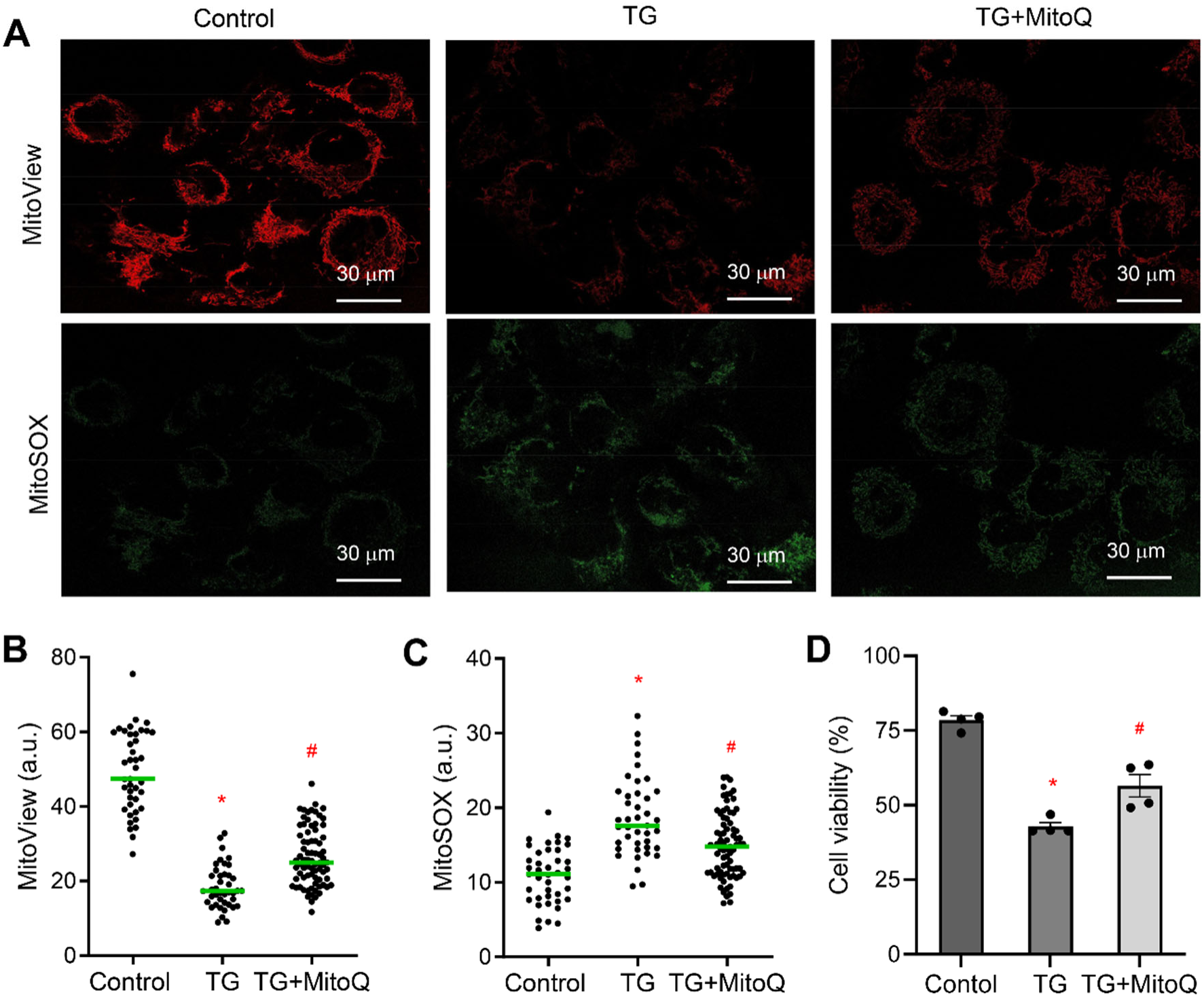
Panx2 knockdown enhances ER stress-associated mitochondrial dysfunction in AC16 cardiomyocytes. (A) Representative confocal images of MitoView for mitochondrial membrane potential (ΔΨ_m_) and MitoSOX for ROS in Panx2 siRNA-transfected AC16 cells under control conditions and following TG (250 nM, 6 h) treatment, with or without MitoQ. (B-D) Quantification of ΔΨ_m_ (B), ROS levels (C) and cell viability (D) in Panx2 siRNA-transfected cells following TG treatment, in the presence or absence of MitoQ. Data are presented as mean ± SEM. Statistical significance was determined using two-way ANOVA test. *: P < 0.05 vs. control. #: P < 0.05 vs. TG. Up to 40 cells from 4 different cultures per group were examined.

To determine the functional contribution of individual UPR branches to the exacerbated cytotoxicity in TG-treated Panx2-deficient cells, siRNA-treated AC16 cells were exposed to TG in the presence or absence of selective UPR pathway inhibitors. PERK inhibition (GSK2656157, 0.3 mM) or IRE1a inhibition (MKC-3946, 10 mM) significantly improved cell viability, with PERK blockade providing the strongest rescue (Figure 3D). In contrast, ATF6a inhibition (Ceapin-A7, 1 mM) had minimal effect (Figure 3C). LDH release assays yielded consistent results, with PERK and IRE1a inhibition attenuating TG-induced cytotoxicity, whereas ATF6a blockade had little impact (Figure 3E). These data support a predominant contribution of PERK signaling, with a contributory role for IRE1a, to the heightened TG sensitivity caused by Panx2 loss.

### Panx2 loss exacerbates ER stress-mediated mitochondrial dysfunction in cardiomyocytes

Given the heightened ER stress signaling and intracellular Ca^2+^ dysregulation observed in Panx2-deficient cells, we next examined whether loss of Panx2 exacerbates ER stress-induced mitochondrial dysfunction. Mock- and Panx2 siRNA-transfected AC16 cells were exposed to TG for 6 h, followed by assessment of mitochondrial membrane potential (ΔΨ_m_) and reactive oxygen species (ROS). In mock-transfected cells, TG significantly reduced ΔΨ_m_ and increased mitochondrial ROS levels (supplemental Figure S1), indicating mitochondrial impairment induced by prolonged ER stress. Notably, Panx2-deficient cells displayed more pronounced mitochondrial depolarization and greater ROS accumulation following TG exposure compared with mock controls (Figure 4A-C). To assess the contribution of mitochondrial oxidative stress to Panx2 deficiency-associated cytotoxicity, we next evaluated the effects of the mitochondria-targeted antioxidant MitoQ^37^ during TG exposure. MitoQ partially restored ΔΨ_m_ and markedly reduced mitochondrial ROS levels (Figure 4A-C). Despite these mitochondrial improvements, MitoQ only modestly increased cell viability, with values markedly below control levels (Figure 4D), indicating that mitochondrial oxidative stress contributes but is not solely responsible for Panx2 deficiency-associated injury.

### Panx2 deficiency lowers the threshold for inflammasome activation and pyroptotic signaling

To define the cell death mechanisms underlying exacerbated cytotoxicity in Panx2-deficient cells, markers of apoptotic cell death were examined in mock- and Panx2 siRNA-treated AC16 cells. Under basal conditions, levels of cleaved caspase-3 (cCasp3) and cleaved PARP (cPARP) were comparable between groups (Figure 5). Following TG exposure, both markers increased in mock- and Panx2-deficient cells, with cPARP levels significantly higher in Panx2 knockdown cells, whereas cCasp3 levels were not proportionally increased, suggesting additional non-apoptotic death mechanisms.

**Figure 5:**
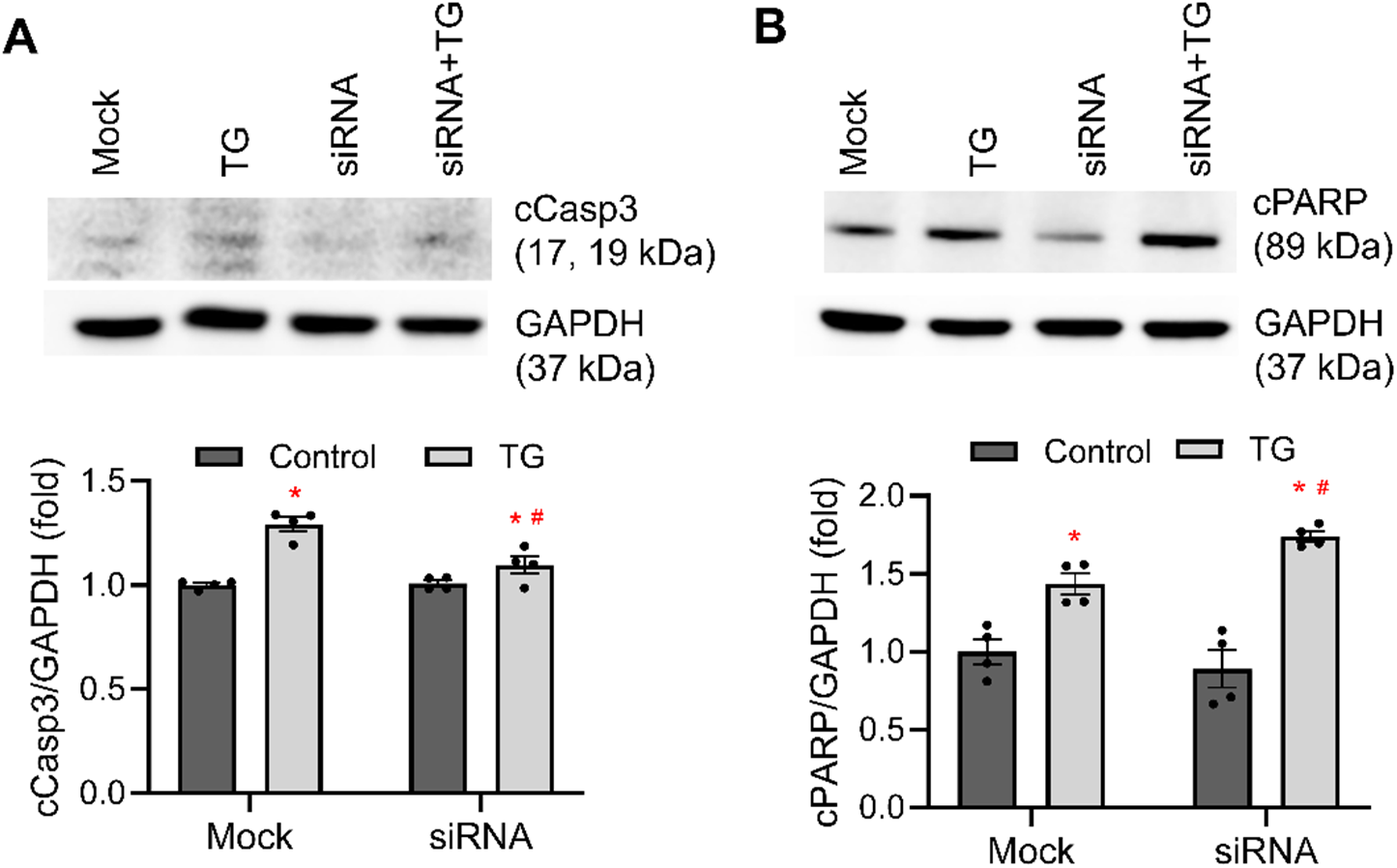
Panx2 deficiency sensitizes cardiomyocytes to ER stress-associated apoptotic cell death. Representative immunoblots and quantification of apoptotic markers cleaved caspase 3 (cCasp3) (A) and PARP (B) in mock- and Panx2 siRNA-transfected AC16 cells following TG exposure (250 nM). Data are presented as mean ± SEM. Statistical significance was determined using two-way ANOVA. *: P < 0.05 vs. control. #: P < 0.05 vs. mock. n=4/group.

We therefore investigated inflammasome-associated signaling and cell death pathways. Mock- and Panx2 siRNA-treated AC16 cells were analyzed under basal conditions and following TG exposure using a time-course immunoblotting approach. Panx2-deficient cells exhibited elevated basal levels of cleaved caspase-1 (cCasp1) compared with mock controls (Figure 6A). Upon TG exposure, cCasp1 levels increased rapidly in Panx2 knockdown cells, peaking between 30-60 minutes, whereas changes in mock-transfected cells were much less and delayed. NLRP3 abundance was also higher at baseline in Panx2-deficient cells and increased further following TG, whereas induction was modest in controls (Figure 6B). Consistent with pyroptotic execution, TG induced gasdermin D (GSDMD) cleavage preferentially in Panx2-deficient cells (Figure 6C). Total pro-caspase-1 and total GSDMD levels were not significantly altered (supplemental Figure S2). Together, these results indicate that Panx2 deficiency sensitizes cardiomyocytes to ER stress-induced inflammasome engagement and pyroptotic signaling.

**Figure 6.**
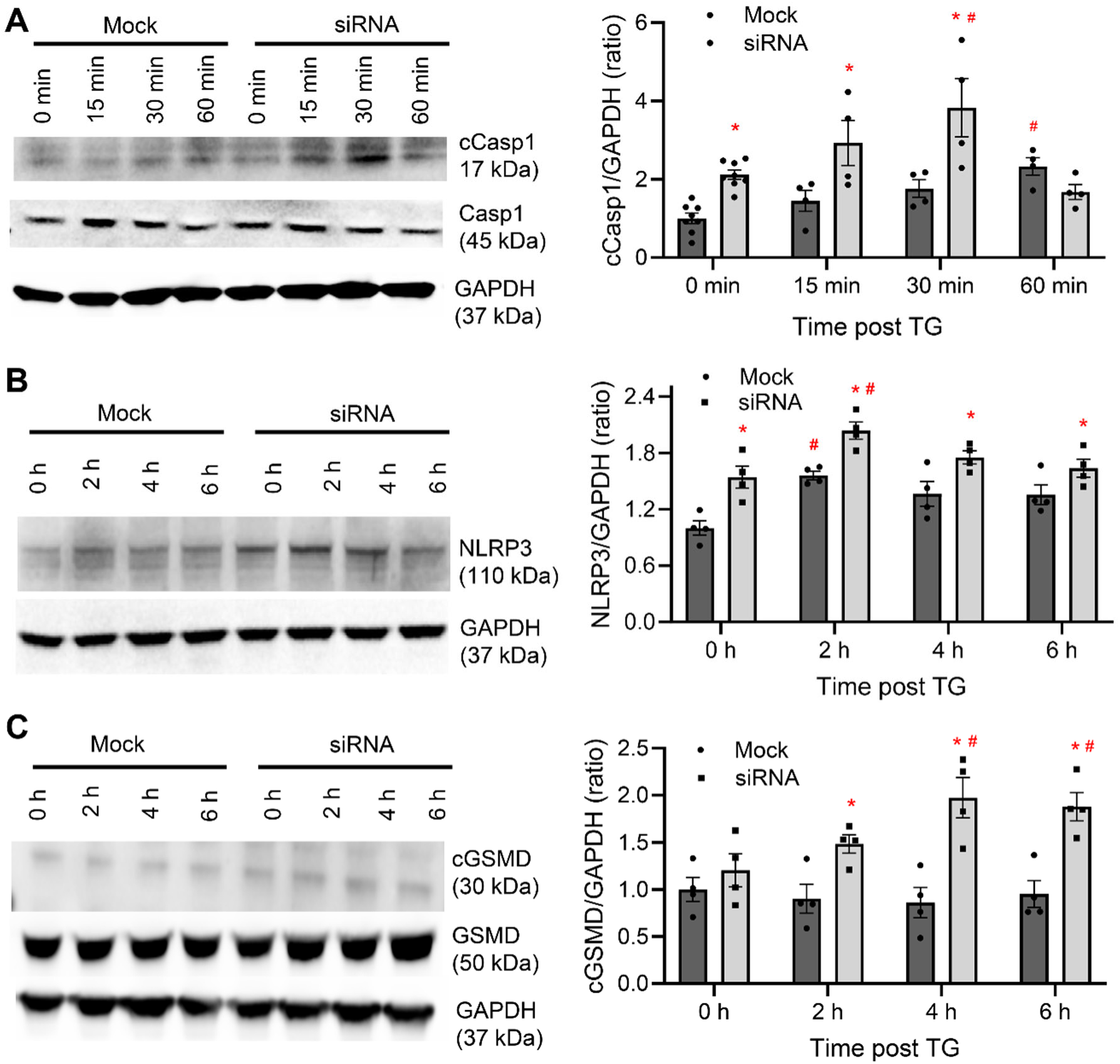
Panx2 deficiency promotes inflammasome activation in cardiomyocytes under ER stress. A-C: Immunoblot analysis and quantification of cleaved caspase-1 (cCasp1) (A), NLRP3 protein levels (B) and cleaved gasdermin D (cGSDMD) (C) in mock- and Panx2 siRNA-transfected AC16 cells under basal conditions and following TG (250 nM) treatment for the indicated times. Protein levels were quantified by densitometry and normalized to GAPDH. Data are presented as mean ± SEM. Statistical significance was determined using two-way ANOVA test. *: P < 0.05 vs. mock. #: P < 0.05 vs. baseline (0 minute). n = 4/group/time point.

### Panx2 overexpression protects cardiomyocyte against ER stress-induced injury via selective suppression of PERK

To determine whether Panx2 overexpression confers protection against ER stress, AC16 cells were transduced with Panx2 adenovirus, resulting in a ∼55-fold increase in Panx2 mRNA and a ∼15-fold increase in protein expression (Figure 7A-B). Cells were then exposed to high-dose TG (1.5 µM) for 24 hours. TG induced substantial cytotoxicity in control cells, whereas Panx2 overexpression significantly reduced LDH release and significantly improved cell viability (Figure 7C-D). Mechanistically, TG-induced PERK activation was nearly abolished by Panx2 overexpression, with p-PERK levels reduced to values comparable to those observed in untreated control cells (Figure 7E). However, p-IRE1α levels were not significantly altered (Figure 7F).

**Figure 7.**
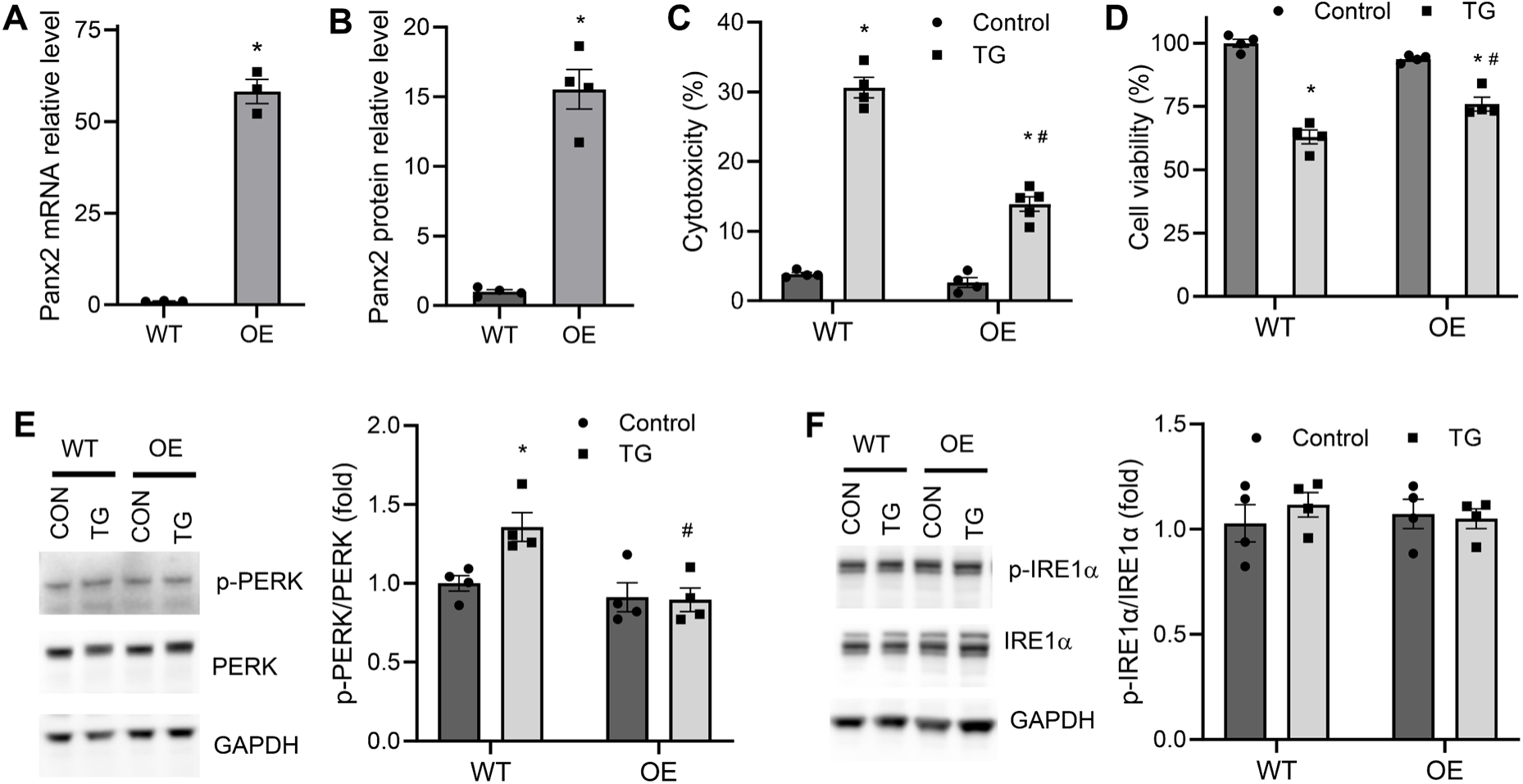
Panx2 overexpression protects cardiomyocytes against ER stress-induced injury. (A) Panx2 mRNA expression measured by RT-qPCR in AC16 cells transduced with control or Panx2 adenovirus. (B) Representative immunoblots and quantification of Panx2 protein levels. (C) LDH release following TG (1.5 µM, 24 h) treatment in wildtype (WT) and Panx2-overexpressing cells. (D) Cell viability or metabolic activity measured under the same conditions as in (C). (E-F) Immunoblot analysis and quantification of phosphorylated PERK (p-PERK) (E) and IRE1a (p-IRE1a) (F) following TG treatment. Data are presented as mean ± SEM. Statistical analysis was performed using two-way ANOVA test. *: P < 0.05 vs. control. #: P < 0.05 vs. WT. n = 4/group.

### Panx2 ablation increases ER stress susceptibility in adult cardiomyocytes

Finally, to validate these findings in primary cells, adult ventricular myocytes were isolated from WT and Panx2^⁻/⁻^ mice. RT-qPCR confirmed near-complete loss of Panx2 transcript expression in Panx2^⁻/⁻^ cardiomyocytes (Figure 8A). Prolonged TG exposure (3 mM, 24 hours) induced greater LDH release in Panx2^⁻/⁻^ cells compared with WT controls (Figure 8B). Calcein AM/EthD-1 staining showed significantly higher cell death in Panx2^⁻/⁻^ cardiomyocytes following TG treatment (supplemental Figure 3, Figure 8C). These data demonstrate that Panx2 loss markedly increases susceptibility of adult cardiomyocytes to ER stress-induced injury.

**Figure 8.**
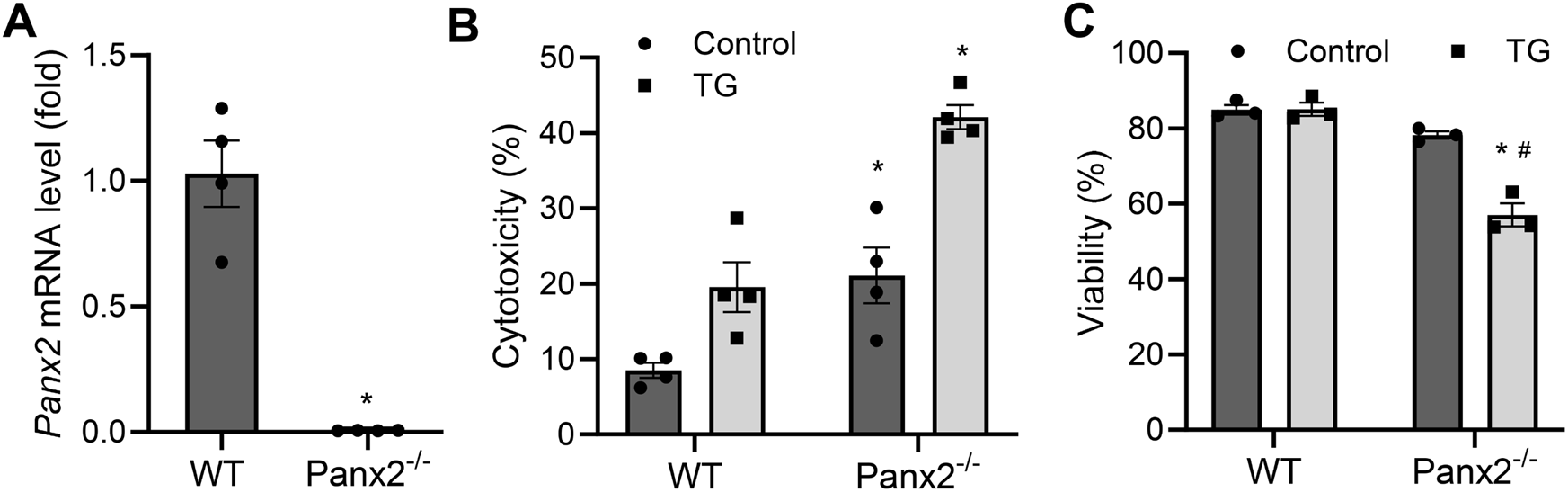
Panx2 ablation increases adult cardiomyocyte susceptibility to ER stress. (A) Panx2 mRNA expression in adult ventricular cardiomyocytes isolated from wildtype (WT) and Panx2^−/−^mice, measured by RT-qPCR. (B) LDH release following TG (3 µM, 24 h) treatment in WT and Panx2^−/−^ cardiomyocytes. (C) Viability of WT and Panx2^−/−^ cardiomyocytes following TG treatment measured by Calcein AM/EthD-1 staining. Data are presented as mean ± SEM. Statistical analysis was performed using two-way ANOVA test. *: P < 0.05 vs. WT. #: P < 0.05 vs. control. n = 4/group.

## Discussion

A central unresolved question in cardiac stress biology is how cardiomyocytes maintain ER Ca^2+^ homeostasis and prevent ER stress from escalating into mitochondrial failure and regulated inflammatory cell death. Although ER stress/UPR activation and disrupted ER-mitochondria Ca^2+^ handling are widely implicated in cardiomyocyte loss across cardiac pathologies, endogenous organelle-localized regulators that set the threshold for maladaptive stress signaling and inflammasome engagement remain incompletely defined. In the present study, we address this gap by identifying Panx2 as an intracellular checkpoint that protects cardiomyocytes from ER stress-driven Ca^2+^ dyeregulation and inflammatory death programs.

Using high-resolution confocal imaging in human AC16 cardiomyocytes, we show that Panx2 does not localize to the plasma membrane but instead exhibits robust enrichment at the ER and close proximity to mitochondria, consistent with its reported distribution in non-cardiac cells^16, 25, 38^. By positioning Panx2 as an organelle-associated intracellular protein, our findings extend the understanding of Panx2 biology to cardiomyocytes and underscore its functional distinction from Panx1, which is exclusively expressed in the plasma membrane and mediates extracellular purinergic signaling through ATP release^39^. The enrichment of Panx2 at ER-mitochondria interfaces is of particular relevance in cardiomyocytes, in which MAMs reside adjacent to key Ca^2+^ regulatory machinery, including IP_3_ receptors and RyR2, and function as hubs for inter-organellar Ca^2+^ and ROS signaling^40^, mitochondrial bioenergetics^41^, and UPR regulation^42^. Our data indicate that Panx2 is not merely spatially localized to these microdomains but functionally constrains Ca^2+^ homeostasis within them. Loss of Panx2 is associated with reduced ER Ca^2+^ stores and elevated basal cytosolic and mitochondrial Ca^2+^ levels, indicating destabilization of ER Ca^2+^ retention and dysregulation of intracellular Ca^2+^ handling. Under ER stress, this Ca^2+^ mishandling is further amplified, resulting in mitochondrial Ca^2+^ overload, oxidative stress, apoptotic cell death and premature engagement of maladaptive UPR signaling. Although further studies are required to fully elucidate the precise molecular mechanisms underlying its action, our findings suggest that Panx2 plays critical roles in maintaining ER-mitochondrial Ca^2+^ homeostasis and cardiomyocyte resilience under stress conditions.

Disruption of ER Ca^2+^ homeostasis is a potent trigger of persistent ER stress^43^. In Panx2-deficient cardiomyocytes, TG exposure markedly enhanced phosphorylation of PERK and IRE1a and cleavage of ATF6a, with all three UPR branches showing elevated basal activation, suggesting that Panx2 helps maintain ER proteostasis even under physiological conditions. Among the three UPR sensors, PERK exhibited the most pronounced and functionally consequential activation, consistent with its early role in eIF2a-mediated translational suppression^44^ and metabolic attenuation^45^. This likely underlies the reduced MTT signal observed at baseline in Panx2-deficient cells, reflecting impaired metabolic capacity rather than overt membrane-compromising cytotoxicity. Enhanced IRE1a and ATF6a activations further indicate unresolved low-grade ER stress across multiple UPR sensors, lowering the threshold for maladaptive signaling upon challenge. It is worth noting that although PERK activation was the predominant determinant of cell injury under Panx2-deficient conditions, partial protection afforded by IRE1a inhibition indicates that IRE1a-dependent signaling also contributes to ER stress-induced cytotoxicity, likely through amplification of downstream inflammatory and death pathways (e.g., JNK/NF-kB-linked programs^46, 47^). Notably, IRE1a can modulate PERK signaling amplitude and expression^48, 49^, and conversely, PERK activity can influence IRE1a signaling^49^, establishing bidirectional crosstalk between these two stress sensors. On the contrastory, while the PERK pathway is required for the proper activation of ATF6a, ATF6a does not directly regulate PERK. Additionally, ATF6 activation primarily functions as a downstream adaptive and cytoprotective transcription factor, not a direct driver of inflammatory cell death. Thus, inhibition of ATF6a fails to blunt cytotoxicity in Panx2-deficient cells once PERK and IRE1a-dependent stress signaling has been engaged. Together, these findings identify Panx2 as a critical regulator of ER stress resolution in cardiomyocytes, with its loss shifting the balance of UPR signaling toward sustained PERK dominance accompanied by secondary IRE1a engagement, thereby promoting a maladaptive, injury-prone stress response.

Although TG classically induces apoptosis or necroptosis^50–52^, severe ER stress can engage non-apoptotic pathways, including caspase-independent death (e.g., autosis)^53^ and inflammatory cell death^54^. In our studies, mitochondrial ROS scavenger MitoQ conferred limited protection, and immunoblotting revealed dissociation between PARP cleavage and caspase 3 activation, suggestive of alternative death modalities^55^. Sustained PERK and IRE1a signaling can converge on NF-κB signaling, priming inflammasome assembly *via* upregulation of NLRP3 and pro-caspase 1^56–58^. In line with this framework, Panx2-deficient cardiomyocytes exhibited pronounced inflammasome activation, characterized by elevated basal levels of cleaved caspase-1 and NLRP3 that were further amplified upon TG exposure, accompanied by robust cleavage of GSDMD. These molecular signatures demonstrate activation of the canonical NLRP3-caspase-1-GSDMD pyroptotic pathway. Mechanistically, the heightened inflammasome responsiveness in Panx2-deficient cells is likely driven by the combined effects of disrupted Ca^2+^ homeostasis and oxidative stress, established upstream triggers of NLRP3 activation^40, 59–61^.

Notably, both apoptotic and pyroptotic cell death markers were detected following TG, indicating that Panx2 loss broadens the spectrum of stress-responsive death programs. The increased vulnerability observed in Panx2-deficient AC16 cells was recapitulated in adult ventricular myocytes from Panx2^−/−^ mice, supporting physiological relevance. Together with prior context-dependent roles of Panx2 in other cell types, the data support a model in which Panx2 function is governed by its intracellular localization and organelle-specific environment rather than plasma-membrane ATP release.

A limitation of the present study is the primary use of AC16 cells, which, despite being a widely used human cardiomyocyte model, do not fully recapitulate the maturation, metabolic, or electrophysiological properties of adult cardiomyocytes^62–64^. While primary adult Panx2^−/−^ mouse data strengthen the conclusions, future validation *in vivo* using cardiac-specific Panx2 models and advanced *ex vivo* systems (e.g., human iPSC-derived cardiac organoids) will be essential for translational advancement. The precise molecular action of Panx2, whether as an ER/MAM channel, regulator of ER/SR Ca^2+^ handling proteins, or modulator of MAM architecture, also requires further investigations.

In summary, this study identifies Panx2 as a novel intracellular regulator of cardiomyocyte stress responses, enriched at ER and MAM interfaces, where it stabilizes Ca^2+^ homeostasis, restrains maladaptive UPR signaling, and limits inflammasome-mediated inflammatory death. These results expand current understanding of pannexin biology by highlighting a role for Panx2 that is distinct from plasma-membrane ATP release and instead centers on organelle-level stress regulation. Given the central involvement of Ca²⁺ dysregulation, mitochondrial dysfunction, and inflammatory cell death in cardiac pathology, Panx2 may represent a promising target for therapeutic strategies aimed at preserving cardiomyocyte resilience by restoring organelle-level homeostasis.

## Data availability

All data generated or analyzed during this study are included in this article and the supporting information files. Any additional data and original data presented in this article are available from the corresponding author upon request.

## Non-standard Abbreviations and Acronyms

ER: endoplasmic reticulum
MAM: mitochondria-associated membrane
MTT: 3-(4, 5-dimethylthiazolyl-2)-2, 5-diphenyltetrazolium bromide
LDH: lactate dehydrogenase
PVDF: polyvinylidene fluoride
SDS-PAGE: sodium dodecyl sulfate-polyacrylamide gel electrophoresis
DMSO: dimethyl sulfoxide
WGA: wheat germ agglutinin
SERCA: sarco/endoplasmic reticulum Ca^2+^-ATPase
UPR: unfolded protein response
PERK: protein kinase RNA-like ER kinase
IRE: iron-responsive element
ATF6: activating transcription factor 6
MitoQ: mitoquinone mesylate
PARP: poly(ADP-ribose) polymerase
NLRP3: nucleotide-binding domain, leucine-rich–containing family, pyrin domain–containing-3
GSDMD: gasdermin D
RyR2: ryanodine receptor 2

